# C/EBPβ-LIP dually activates glycolysis and the malate aspartate shuttle to maintain NADH/NAD+ homeostasis

**DOI:** 10.1101/2020.10.09.333104

**Authors:** Tobias Ackermann, Hidde R. Zuidhof, Gertrud Kortman, Martijn G. S. Rutten, Mathilde Broekhuis, Mohamad Amr Zaini, Götz Hartleben, Cornelis F. Calkhoven

## Abstract

Oncogene-induced metabolic reprograming supports cell growth and proliferation. Yet, it also links cancer cell survival to certain metabolic pathways and nutrients. In order to synthesise amino acids and nucleotides *de novo* for growth and proliferation, cancer cells depend on glycolysis, the cytoplasmic oxidation of glucose, which generates necessary metabolic intermediates and ATP. During glycolysis, NAD+ is used as the oxidizing agent and is thereby reduced into NADH. To ensure high glycolysis rates and maintain NADH/NAD+ homeostasis, cytoplasmic NAD+ has to be regenerated. The mitochondria are the major sites of NADH reoxidation into NAD+ where NADH-derived electrons enter the electron transport chain for ATP production. Since NADH/NAD+ cannot cross membranes, the malate-aspartate shuttle (MAS) or the glycerol-3-phosphate shuttle (GPS) are used as intermediate electron carriers. In addition, cytoplasmic NAD+ is generated by NADH-electron transfer to pyruvate, reducing it to lactate (the Warburg effect). NADH/NAD+ homeostasis plays a pivotal role in cancer cell survival, but our knowledge about the involved regulatory mechanisms is still limited. Here, we show that the proto-oncogenic transcription factor C/EBPβ-LIP stimulates both glycolysis and the MAS. Inhibition of glycolysis with ongoing C/EBPβ-LIP-induced MAS activity results in NADH depletion and apoptosis that can be rescued by inhibiting either the MAS or other NADH-consuming processes. Therefore, beyond the discovery of C/EBPβ-LIP as a dual activator of glycolysis and the MAS, this study indicates that simultaneous inhibition of glycolysis and lowering of the NADH/NAD+ ratio may be considered to treat cancer.

## Introduction

Cancer cells reprogram their metabolism to support *de novo* synthesis of macromolecules that are needed for cell growth and proliferation (Hanahan and Weinberg, 2011). Most prominently, cancer cells increase glucose uptake and metabolise glucose by aerobic glycolysis, which was first recognised by Otto Warburg (reviewed in (Koppenol et al., 2011)). Later, it was shown that cancer cells maintain high rates of both glycolysis and oxidative phosphorylation (OXPHOS) to meet the high demand of energy and substrates for anabolic processes (DeBerardinis and Chandel, 2016; Vander Heiden and DeBerardinis, 2017). The high glycolytic flux provides the cancer cell with macromolecules by uncoupling glycolysis from the mitochondrial tricarboxylic acid (TCA) cycle and diverting glucose carbon into biosynthetic pathways, including the pentose phosphate pathway, hexosamine pathway and serine biosynthetic pathway (Pavlova and Thompson, 2016). During glycolysis and serine biosynthesis NAD+ serves as an electron acceptor, for reactions catalysed by glyceraldehyde-3-phospate dehydrogenase (GAPDH) and phosphoglycerate dehydrogenase (PHGDH), and is reduced to NADH. Consequently, the NADH/NAD+ ratio in cancer cells is usually very high (Hung et al., 2011). To sustain a high glycolytic flux and allow serine biosynthesis, NAD+ has to be regenerated. The mitochondria are major sites of NAD+-regeneration by complex I of the electron transport chain (ETC), where NADH-derived electrons contribute to oxidative respiration and ATP production (Gui et al., 2016). Yet, NAD+ and NADH cannot pass the mitochondrial membranes and therefore cells use substrate cycles that transport the electrons derived from oxidation of cytosolic NADH into the mitochondria. One such cycle is the malate-aspartate shuttle (MAS), which uses malate as intermediate electron carrier to transport the electrons from cytosolic NADH to mitochondrial NADH (Borst, 2020). The other cycle is the glycerol-phosphate shuttle (GPS), which uses Glycerol-3-phosphate as intermediate to transport the electrons over the membrane to mitochondrial FADH that enters the ETC at complex II (**Supplementary Fig. 1A**). In either way the carbon flux of the glycolysis is coupled to mitochondrial function (Birsoy et al., 2015; DeBerardinis et al., 2008). There are early reports that cancer cells depend on the MAS for biomolecule synthesis and proliferation, while this is less clear for the GPS (Greenhouse and Lehninger, 1976; Hanse et al., 2017). When mitochondrial NAD+ regeneration does not suffice, which is mostly the case for the high glycolytic flux in cancer cells, cytosolic NAD+ is regenerated in addition through reduction of pyruvate into lactate by the enzyme lactate dehydrogenase (LDH) (the Warburg effect). Yet, LDH activity alone cannot provide sufficient cytosolic NAD+, and through lactate secretion glucose carbon is lost for biomolecule synthesis (DeBerardinis et al., 2008).

The transcription factor CCAAT/enhancer binding protein beta (C/EBPβ) is known to regulate organismal metabolism (Millward et al., 2007; Zidek et al., 2015). The *Cebpb*-mRNA is translated into three protein isoforms; two transcriptional activators C/EBPβ-LAP1 and - LAP2 (also named LAP* and LAP) and the N-terminally truncated protein isoform and transcriptional inhibitor C/EBPβ-LIP (hereinafter referred to as LAP and LIP)(Calkhoven et al., 2000; Descombes and Schibler, 1991). LIP expression is tightly controlled by the mTORC1-4E-BP pathway and involves a *cis*-regulatory upstream open reading frame (uORF) in the *Cebpb*-mRNA leader sequence (Calkhoven et al., 1994; Calkhoven et al., 2000). Overexpression of LIP induces cellular transformation (Calkhoven et al., 2000) and increases tumour incidence in mice (Begay et al., 2015), while LIP deficiency reduces tumour incidence in mice (Müller et al., 2018). Moreover, LIP is highly expressed in breast cancer, ovarian cancer, colorectal cancer and anaplastic large cell lymphoma (Jundt et al., 2005; Quintanilla-Martinez et al., 2006; Rask et al., 2000; Sundfeldt et al., 1999; Zahnow et al., 2001; Zahnow et al., 1997). Recently, we demonstrated that LIP induces cancer metabolism with increased glycolysis and mitochondrial respiration through regulation of the let-7/LIN28B circuit (Ackermann et al., 2019).

Here, we show that the simultaneous stimulation of glycolysis and the MAS in high LIP expressing cells causes reliance on glycolysis for NADH/NAD+ homeostasis and cell viability. Inhibition of glycolysis in cells with high LIP expression results in depletion of NADH and low NADH/NAD+ ratios, because the MAS continues to oxidise cytosolic NADH into NAD+ and the mitochondrial NADH is used for ATP production. This condition is associated with apoptosis, which can be rescued by inhibition of the MAS or other NADH-oxidising processes. Our data indicate that low NADH/NAD+ ratios are part of the 2-DG mediated toxicity and that metabolic reprogramming by LIP or through otherwise increased MAS activity makes cancer cells vulnerable to glycolytic inhibitors.

## Results

### LIP renders cells dependent on glucose metabolism

We examined the C/EBPβ isoform specific effects on cellular metabolism by measuring the extracellular acidification rate (ECAR) as an indicator for glycolytic flux, and the oxygen consumption rate (OCR) as a measure for mitochondrial metabolism (Seahorse XF96). Similar to what we showed earlier (Ackermann et al., 2019), experimental expression of LIP in immortalised *Cebpb*-knockout MEFs increased both basal and maximal ECAR, while expression of LAP had no effect on the ECAR (**Fig. 1A**). The LIP-induced increase in ECAR was abrogated by treatment with 2-deoxyglucose (2-DG), which inhibits glycolysis (**Supplementary Fig. 1B**), Together these data show that LIP enhances the glycolytic flux. In addition, expression of LIP and to a lesser extent LAP increased the OCR in the cells (**Fig. 1A and Supplementary Fig. 1B**) indicating that both LIP and LAP stimulate mitochondrial metabolism with different potential. Next, we investigated whether LIP-expressing cells depend on the increased glycolytic flux for cell proliferation and survival. Withdrawal of glucose resulted in a strong decrease in viable LIP-expressing cells after three days of culture compared to cells cultured in the presence of glucose (25 mM) (**Fig. 1B**). Glucose deprivation did not result in altered cell numbers for LAP-expressing cells or empty vector (EV) control cells compared to glucose containing media (**Fig. 1B**). Deprivation of the alternative carbon source glutamine alone or together with glucose resulted in a strong decrease in cell numbers in all three cell lines (EV, LIP, LAP), confirming a general requirement of glutamine for cell proliferation independent of LIP or LAP expression (**Fig. 1B**). The induction of Caspase 3/7 activity in LIP-but not in LAP-expressing cells showed that only LIP-expressing cells undergo apoptosis upon glucose starvation (**Fig. 1C**). In line with the glucose-deprivation experiments, treatment with the glycolytic inhibitor 2-DG abrogates LIP-induced increase in ECAR (**Supplementary Fig. 1B**) and drives LIP-expressing cells into apoptosis as was shown by a strong increase in Caspase 3/7 activity (**Fig. 1D**) and microscopy (**Fig. 1E**), while LAP-expressing and control (EV) cells survive under this condition. Hence, these data show that proliferation and survival of LIP-expressing MEFs highly depends on glucose metabolism.

**Figure 1.**
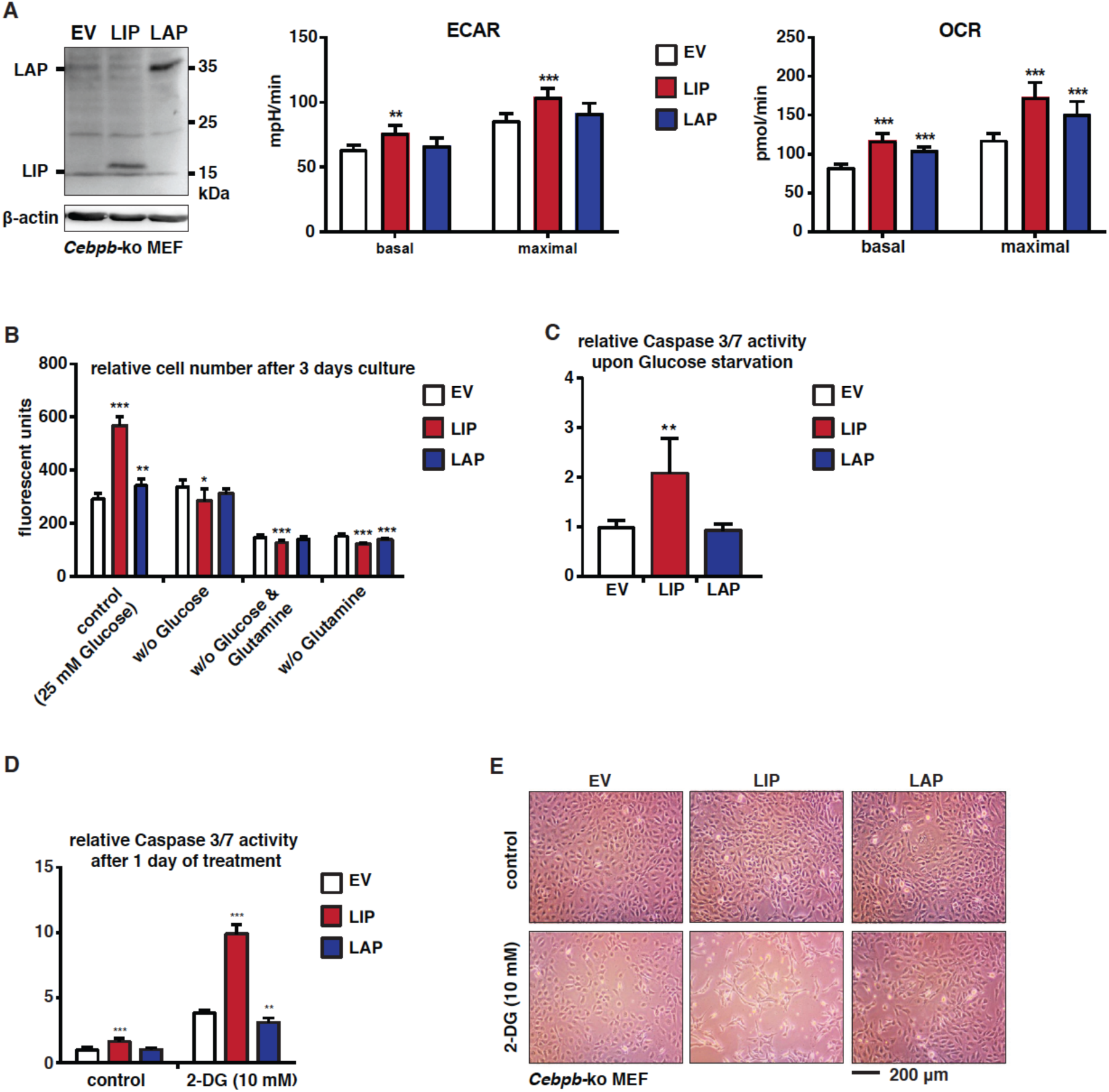
LIP induces reliance on glycolysis for cell proliferation and survival. (**A**) The immunoblot at the left shows expression of LAP, LIP and β-actin as loading control in *Cebpb*-ko fibroblasts transfected with expression vectors for LAP, LIP or empty vector (EV) control. The bar graphs show values of ECAR and OCR associated with LAP, LIP or EV expression (n=6). (**B**) Relative cell numbers of *Cebpb*-ko fibroblasts transfected with expression vectors for LAP, LIP or empty vector (EV) control after 3 days of culture in media with or without glucose and/or glutamine (n=5). (**C**) Relative Caspase3/7 activity in *Cebpb*-ko fibroblasts transfected with expression vectors for LAP, LIP or empty vector (EV) control after 3 days of culture in medium without glucose (n=5). (**D**) Relative Caspase3/7 activity of *Cebpb*-ko fibroblasts transfected with expression vectors for LAP, LIP or empty vector (EV) control after one day of treatment with 2-DG (n=5). (**E**) Microscopic pictures of *Cebpb*-ko fibroblasts transfected with expression vectors for LAP, LIP or empty vector (EV) control after one day of treatment with 2-DG. Statistical differences were analysed by Student’s t-tests. Error bars represent SD, * p<0.05, ** p<0.01, *** p<0. 001.

As a cellular model for further studies we chose breast cancer cell lines, since high expression levels of LIP have been described for aggressive breast cancer types (Zahnow et al., 1997). Immunoblot analysis of a panel of breast cancer cell lines revealed high endogenous LIP expression (high LIP/LAP ratio) in triple negative breast cancer (TNBC) subtype, while low LIP expression with lower LIP/LAP ratios were found in luminal A subtype breast cancer cell lines (**Fig. 2A**). To examine whether high LIP expression is associated with higher sensitivity to inhibition of glycolysis we generated dose-response curves for 2-DG treatment and cell multiplication. From the TNBC cell lines, the BT20 cells that express the highest LIP levels were most sensitivity to 2-DG (IC50=0.6 mM), followed by BT549 (IC50=2.4 mM) and MDA-MB-231 (IC50=4.1 mM) (**Fig. 2B and Supplementary Fig. 2A**). The luminal A type cells with low LIP expression showed a poorer response to 2-DG treatment (IC50>20 mM) (**Fig. 2B and Supplementary Fig. 2A**). Furthermore, the sensitivity to 2-DG correlated with increased Caspase3/7 activity as a measurement for apoptosis in BT20 cells and to lesser extent in BT549 and MDA-MB-231 cells, while Luminal A breast cancer cells even showed a slight decrease in Caspase3/7 activity in response to 2-DG (**Fig. 2C**). To address whether the high sensitivity for 2-DG of BT20 cells depends on C/EBPβ expression we generated *Cebpb*-ko BT20 cells by CRISPR/Cas9 genome editing (**Fig. S2B**). In three independent knockout clones (**Fig. 2D**) the sensitivity to 2-DG was reduced as measured by cell multiplication (**Fig 2E and Supplementary Fig. 2C**) and by a significant reduction in Caspase 3/7 activation, although to various extents (**Fig. 2F**). Next, we investigated whether overexpression of LIP in the low LIP-expressing T47D and MCF-7 Luminal A cell lines would result in increased sensitivity to 2-DG, as measured by cell multiplication.

**Figure 2.**
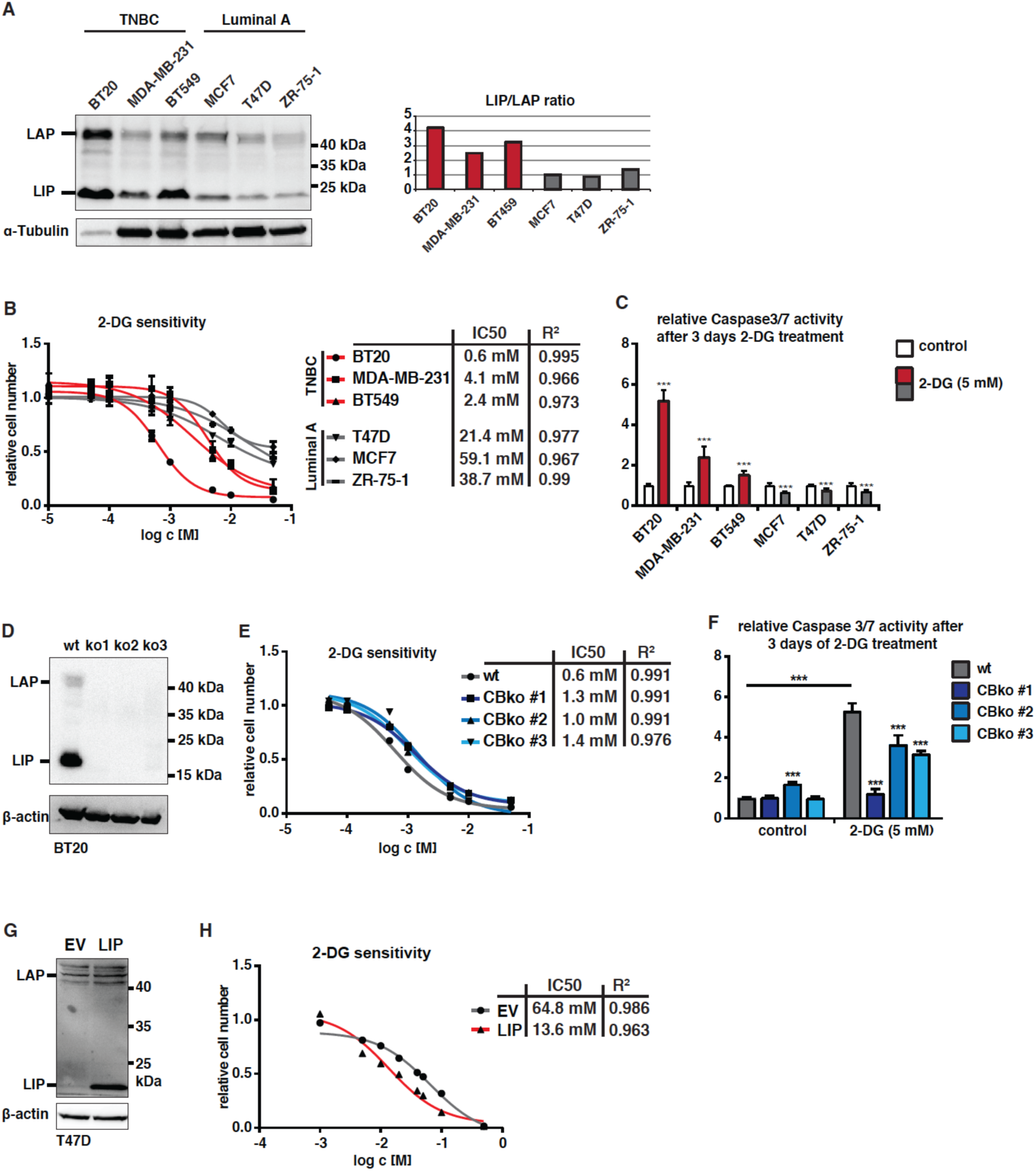
High LIP/LAP ratios render TNBC cell lines sensitive to 2-DG. (**A**) Immunoblot showing expression of LAP, LIP and α-tubulin as loading control in the TNBC BT20, MDA-MB-231, BT549, and Luminal A MCF7, T47D, ZR-75-1 breast cancer cell lines with quantified LIP/LAP ratios at the right. (**B**) Dose-response-curve of the six breast cancer cell lines mentioned in A after 3 days of 2-DG treatment (n=5). (**C**) relative Caspase 3/7 activity of the six breast cancer cell lines mentioned in A after 3 days of 2-DG treatment (n=5). (**D**) Immunoblot showing expression of LAP, LIP and β-actin as loading control in wt BT20 cells and three clones of CRIPR/Cas9 derived *Cebpb*-ko BT20 cells. (**E**) Dose-response-curve of wt BT20 and *Cebpb*-ko BT20 cells after 3 days of 2-DG treatment (n=5). **F**) Relative Caspase 3/7 activity in wt BT20 and *Cebpb*-ko BT20 cells after 3 days of 2-DG treatment (n=5). (**G**) Immunoblot showing expression of LAP, LIP and β-actin as loading control in T47D cells transfected with expression vectors for LIP or empty vector (EV) control. (**H**) Dose-response-curve of T47D cells expressing LIP or EV control and after 3 days of 2-DG treatment (n=5). Statistical differences were analysed by Student’s t-tests. Error bars represent SD, * p<0.05, ** p<0.01, *** p<0. 001. IC50 values of dose-response are determined by R-squared (R^2^) statistical analysis.

The higher LIP expression indeed increased the sensitivity to 2-DG by 5-fold for T47D and 2.3-fold for MCF7 (**Fig. 2G,H and Supplementary Fig. 2D, E**). However, LIP overexpression did not induce Caspase 3/7 activity in T47D cells suggesting that, either in TNBC cells other factors contribute to 2-DG-induced apoptosis in addition to LIP, or that in the Luminal A cells mechanisms are active that prevent apoptosis (**Supplementary Fig. 2F**). Taken together, these data show that high levels of LIP expression render TNBC cells dependent on glycolysis for cell proliferation and survival.

### LIP increases the use of glycolysis-derived NADH for mitochondrial respiration

We next asked why high LIP expression increases the dependence on glycolysis. We first examined whether ATP/ADP ratios change upon LIP-induction, which could be involved in inducing apoptosis (Vander Heiden et al., 2009). One source of ATP production in cancer cells is the high glycolytic flux. Inhibition of glycolysis with 2-DG reduced the ECAR as a measure for glycolytic flux to the same extent in T47D-LIP cells and control T47D cells (**Fig. 3A**). The expectation was that ATP/ADP ratios would decline to comparable degrees. However, compared to the 51% reduction in T47D-EV cells 2-DG treatment more strongly reduced the ATP/ADP ratio in the T47D-LIP cells with 70% (**Fig. 3B**). Thus, in T47D-LIP cells, in addition to the ATP produced during aerobic glycolysis other glucose-dependent pathway(s) contribute for 19% to the production of cellular ATP, which if prevented would compromise cell proliferation and/or viability. We reasoned that a mechanism involving glycolysis-coupled stimulation of mitochondrial respiration and the associated generation of ATP in the electron transport chain (ETC) might explain the extra ATP production induced by LIP.

**Figure 3.**
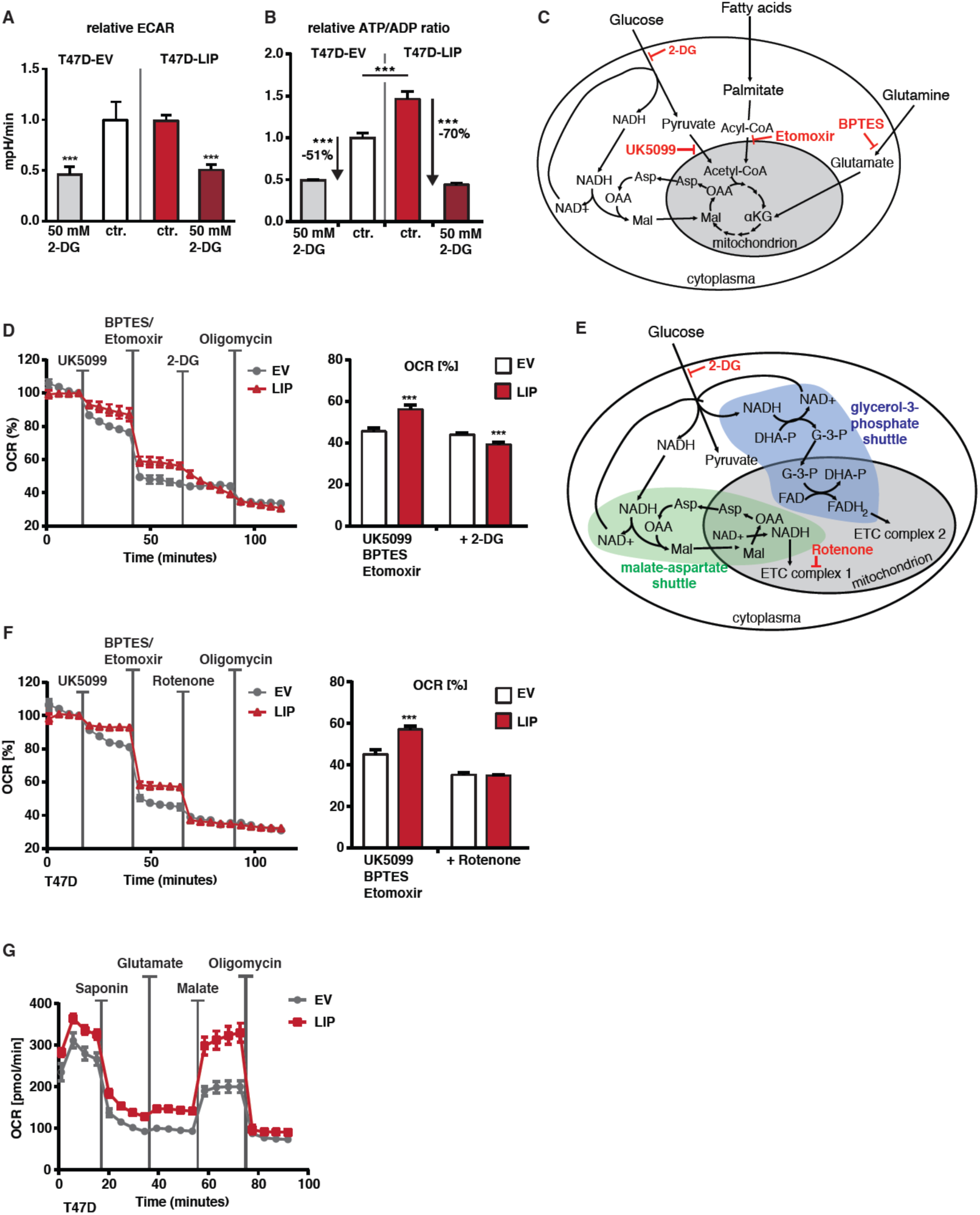
LIP stimulates the MAS and its usage of cytoplasmic NADH. (**A**) Relative ECAR of LIP expressing or control (EV) T47D cells before and 10 minutes after with treatment with 2-DG (n=5). (**B**) Relative ATP/ADP ratios in LIP expressing or control (EV) T47D cells after one day of 2-DG treatment or without treatment (ctr.) (n=5). (**C**) Schematic representation of the flow of metabolites between cytoplasm and the mitochondria and inhibitors of specific pathways. (**D**) OCR of LIP expressing or control (EV) T47D cells with subsequent injection of UK5099, BPTES plus Etomoxir, 2-DG and oligomycin (n=6). Bar graph at the right shows a different representation of the data (n=6). (**E**) Schematic representation of the MAS and the GPS, the flow of involved metabolites and used inhibitors. (**F**) OCR of LIP expressing or control (EV) T47D cells with subsequent injection of UK5099, BPTES plus Etomoxir, rotenone and oligomycin (n=6). Bar graph at the right shows a different representation of the data (n=6). (**G**) OCR of LIP expressing or control (EV) T47D cells with subsequent injection of saponin (permeabilization of cell membrane), glutamate, malate and oligomycin (n=6). Statistical differences were analysed by Student’s t-tests. Error bars represent SD, *** p<0. 001.

Mitochondrial metabolism is fuelled mainly by pyruvate (as a source for Acetyl-CoA) and NADH from glycolysis, Acyl-CoA from fatty acid catabolism, and α-Ketoglutarate from glutaminolysis (**Fig. 3C**). The possible engagement of the individual pathways can be studied by measuring changes in oxygen consumption rate (OCR; Seahorse XF96) upon treatment with specific pathway inhibitors (**Fig. 3D**). Inhibition of the mitochondrial pyruvate carrier with the drug UK5099 reduced the OCR in T47D-LIP cells to a lesser extent compared to control T47D cells, suggesting that pyruvate from glycolysis is not a critical metabolite to fuel respiration in T47D-LIP cells (**Fig. 3C**). When in addition the use of palmitate and glutamine by the mitochondria is blocked with Etomoxir and BPTES (Bis-2-5-phenylacetamido-1,3,4-thiadiazol-2-ylethyl sulphide) the T47D-LIP cells still maintained higher OCR compared to the T47D-EV cells (**Fig. 3D**). Under these conditions the NADH generated by glycolysis can still fuel mitochondrial respiration. Therefore, we used 2-DG to inhibit glycolysis on top of the treatment with UK5099/Etomoxir/BPTES and found the OCR of T47D-LIP cells now decreased to a similar extent than control T47D cells (**Fig. 3D and E**). These data show that LIP stimulates the usage of cytosolic glycolysis-derived NADH for promoting mitochondrial respiration.

### LIP stimulates the malate aspartate shuttle (MAS)

The two pathways that transport electrons derived from cytosolic NADH into the mitochondria are the malate-aspartate shuttle (MAS) and the glycerol-phosphate shuttle (GPS) (**Supplementary Fig. 1A**). Because the MAS generates NADH in the mitochondria that enters complex I of the ETC and the GPS generates FADH that enters the ETC at complex II, we were able to discriminate between those two by inhibition of complex I with rotenone. Following inhibition of mitochondrial pyruvate (UK5099), palmitate (Etomoxir) and glutamine (BPTES) metabolism, rotenone abrogated the difference in OCR caused by LIP expression in T47D cells, indicating that MAS is the critical pathway involved in enhancing mitochondrial respiration (**Fig. 3F**). Next, we examined the capacity of the MAS in T47D-LIP versus control T47D cells by experimentally applying exogenous malate after cell membrane permeabilization using saponin. Permeabilization of the cell membrane resulted in a strong reduction of the OCR for both cell lines, because substrates for mitochondrial respiration are leaking out of the cell (**Fig. 3G**). The subsequent supply of glutamate slightly increased the OCR in both cell lines while the supply of the MAS substrate malate resulted in a strong increase in OCR in the T47D-LIP cells and to a much lesser extent in the control T47D cells (**Fig. 3G**), which was reduced to the same level by treatment with rotenone (**Fig. 3F**). These data show that LIP stimulates the MAS, transporting electrons from cytoplasmic NADH into the mitochondria, to stimulate mitochondrial respiration.

### Altered NADH usage causes apoptosis in LIP overexpressing cells

Next, we asked whether the LIP-induced increase in MAS renders the cells sensitive to inhibition of glycolysis. Hypothetically, inhibition of glycolysis may lead to very low NADH/NAD+ ratios with potentially detrimental effects on cell viability, particularly in cells with high LIP expression because the LIP-driven MAS continues to convert cytosolic NADH into NAD+. Inhibition of glycolysis in the high-LIP BT20 cells with 2-DG resulted in a strong decrease in NADH/NAD+ ratio (**Fig. 4B**) and induction of apoptosis (Caspase 3/7 activity) (**Fig. 4C**). In an attempt to increase the NADH/NAD+ ratio and rescue cell viability, the MAS was inhibited by treatment with aminooxyacetic acid (AOA) that broadly inhibits transaminases including the aspartate aminotransferase of the MAS (**Fig. 4A**). Treatment with AOA fully restored the cellular NADH/NAD ratio in 2-DG treated cells (**Fig. 4B**) and resulted in a significant although not full decrease in apoptosis (**Fig. 4C**). The incomplete rescue from apoptosis may be due to AOA being a “promiscuous” drug, inhibiting all transaminases in the cell, which potentially may affect cell survival in addition (Thornburg et al., 2008). Notably, the ATP/ADP ratio in the 2-DG treated cells did not increase upon AOA treatment (**Fig. 4D**) and therefore cannot contribute to the increase in cell viability. In the TNBC cell lines BT549 and MDA-MB-231 that have lower LIP expression 2-DG treatment likewise lowered the NADH/NAD+, although to a lesser extent then in BT20 cells (**Supplementary Fig. 3A and D**). Also, in these cell lines subsequent inhibition of the MAS with AOA restored the cellular NADH/NAD+ ratio, resulted in a significant decrease in apoptosis (**Supplementary Fig. 3B and E**), and did not alter the ATP/ADP ratio in the 2-DG treated cells (**Supplementary Fig. 3C and F**). Therefore, the data suggest that LIP-stimulated MAS activity depletes NADH from the cytoplasm and when NADH is not continuously replenished by a high glycolytic flux the low NADH/NAD+ ratios compromise cell viability.

**Figure 4.**
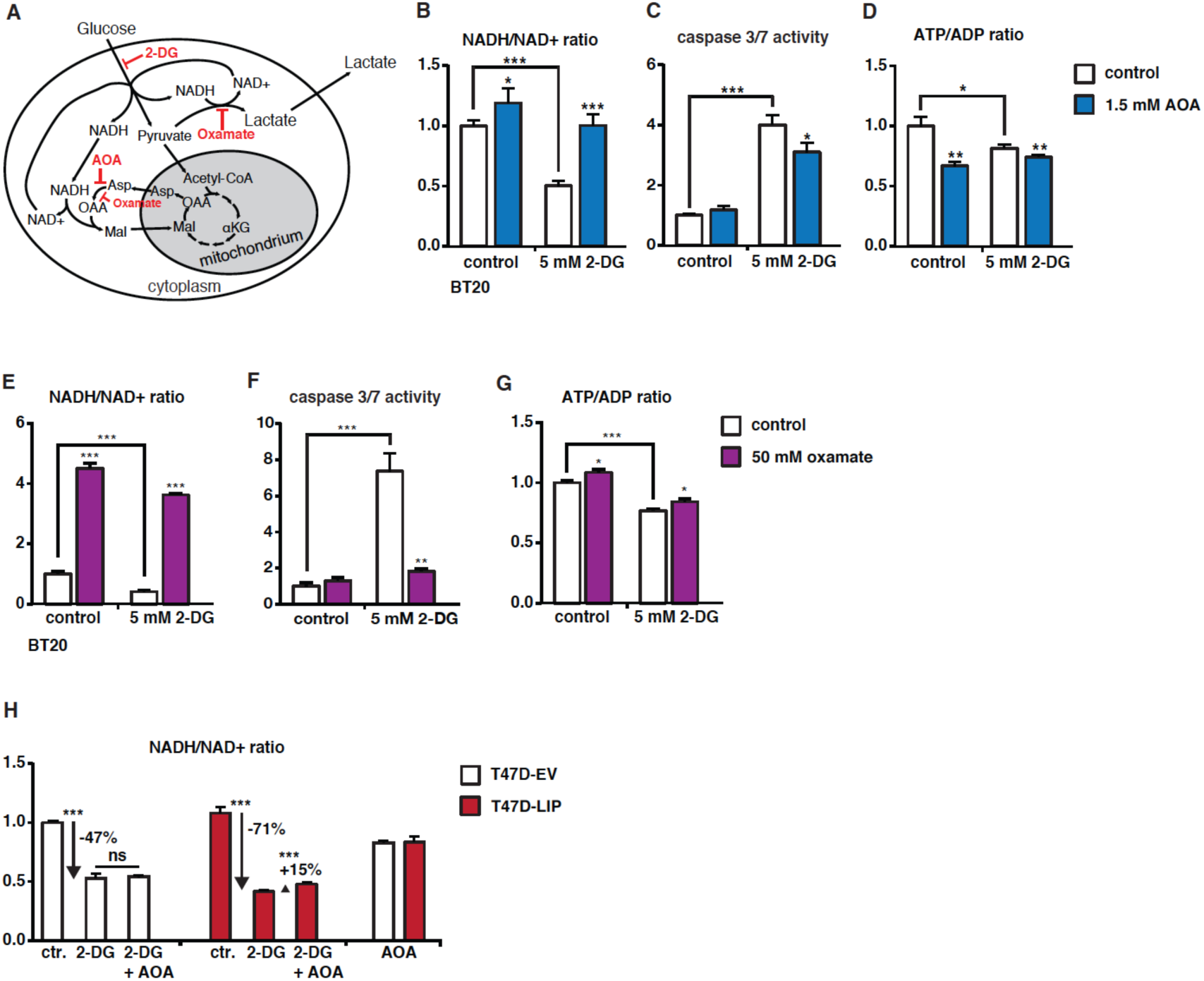
Inhibition of NADH-consuming processes reduces toxicity of 2-DG. (**A**) Schematic representation of cytoplasmic NADH-consuming processes MAS and LDH and their inhibitors AOA and oxamate, respectively, and the glycolytic inhibitor 2-DG. (**B**) Relative NADH/NAD+ ratios in BT20 cells after 1 day of treatment with solvent (control), 2-DG, AOA, or 2-DG plus AOA (n=4). (**C**) Relative caspase3/7 activity of BT20 cells after 3 days of treatment with solvent, 2-DG, AOA or 2-DG plus AOA (n=4). (**D**) Relative ATP/ADP ratios in BT20 cells after 1 day of treatment with solvent, 2-DG, AOA, or 2-DG plus AOA (n=4). (**E**) Relative NADH/NAD+ ratios in BT20 cells after 1 day of treatment with solvent, 2-DG, oxamate or 2-DG plus oxamate (n=4). (**F**) Relative caspase3/7 activity of BT20 cells after 3 days of treatment with solvent, 2-DG, oxamate or 2-DG plus oxamate (n=4). (**G**) Relative ATP/ADP ratios in BT20 cells after 1 day of treatment with solvent, 2-DG, oxamate or 2-DG plus oxamate (n=4). (**H**) Relative NADH/NAD+ ratios in T47D-LIP overexpressing or T47D-EV (empty vector) control cells after treatment of solvent (ctr.), 2-DG or AOA (n=4). Statistical differences were analysed by Student’s t-tests. Error bars represent SD, * p<0.05, ** p<0.01, *** p<0. 001.

Oxamate is a structural analogue of pyruvate that inhibits LDH and in a side reaction also inhibits the MAS enzyme aspartate aminotransferase (AAT) to a lesser extent (**Fig. 4A**) (Rej, 1979; Thornburg et al., 2008). Treatment of BT20 cells with oxamate greatly increased NADH/NAD+ ratios, reflecting the inhibition of the NADH-consuming reactions of LDH and AAT (**Fig. 4E**). When glycolysis and therefore the pyruvate supply for LDH is inhibited by 2-DG, oxamate strongly increased NADH/NAD+ ratios, likely by inhibition of AAT (**Fig. 4E**). Importantly, the strong upregulation of apoptosis measured by caspase 3/7 activity following 2-DG treatment was almost completely reverted by the NADH/NAD+ ratio restoring oxamate treatment (**Fig. 4F**). In this case the ATP/ADP ratio slightly increased in the 2-DG and oxamate treated cells, but did not reach control levels (**Fig. 4G**).

Altogether, these data show that cells do not tolerate inhibition of glycolysis together with active NADH-consuming processes, and that cells can be rescued from apoptosis by restoring NADH/NAD+ ratios through inhibition of NADH-consuming pathways. In further support that high-LIP/high MAS contributes to the toxicity of 2-DG is that treatment with 2-DG resulted in a stronger drop in NADH/NAD+ ratios (−71%) in T47D-LIP cells compared to control T47D cells (−47%) (**Fig. 4H**). Subsequent inhibition of the MAS by AOA restored the NADH/NAD+ ratio with +15% in T47D-LIP but not in T47D-EV cells, showing that LIP overexpression induces the MAS in these luminal A breast cancer cells similar to the high endogenous LIP expression in TNBC cell lines.

## Discussion

The data presented in this study suggest that C/EBPβ-LIP not only induces glycolysis but in addition secures continuation of the glycolytic flux by stimulating the regeneration of cytoplasmic NAD+ through the MAS. The coordination of these two interdependent processes that are particularly important for cancer cell growth and proliferation has, to our knowledge, thus far not been shown for other oncogenic factors. Upon glucose withdrawal or inhibition of glycolysis with 2-DG the ongoing LIP-induced MAS strongly decreases the cellular NADH/NAD+ ratio, inducing apoptosis. In cells with high endogenous levels of LIP the sensitivity to 2-DG can be diminished by depletion of C/EBPβ, inhibition of the MAS or other NADH-oxidising processes. Hence, LIP may mark cancer cells for sensitivity to therapeutic strategies targeting glycolysis (e.g. 2-DG).

The MAS is a key process in the cell to connect metabolic pathways in the mitochondria and the cytoplasm and has been linked to cancer metabolism (Borst, 2020), for example in pancreatic cancer or non-small lung cancer where amplification of Malate dehydrogenase 1 (MDH1) was detected (Wang et al., 2016). MDH1 is the key enzyme in the MAS that catalyses NADH/NAD+-dependent reversible oxidation of malate into oxaloacetate in the cytosol. In lung cancer cells, the enhanced MAS activity is required as an alternative to LDH-catalysed NAD+ generation, since glucose carbons are shuttled into biosynthetic pathways and therefore the pyruvate supply to LDH is not sufficient to regenerate NAD+ to maintain a high glycolytic flux (Hanse et al., 2017). The MAS allows cancer cells to efficiently use the glycolysis for both energy production and anabolic processes and thereby can stimulate proliferation or survival upon glutamine starvation (Tajan et al., 2018). Our data indicate that LIP induces an increase in MAS activity that may be part of the oncogenic activities of LIP (Ackermann et al., 2019; Begay et al., 2015). Our preliminary experiments on regulation of the key factors SLC25A11 (malate/α-ketoglutarate carrier), SLC25A12 and SLC25A13 (citrin, glutamate/aspartate carrier), MDH1 (malate dehydrogenase, cytoplasmic), MDH2 (mitochondrial), GOT1 (glutamic oxaloacetic aminotransferase, cytoplasmic) and GOT2 (mitochondrial) as well as on the methylation status of MDH1 known to regulate its function did not show LIP-dependent changes. Therefore, further studies will concentrate on discovery of the underlying molecular mechanisms of LIP-MAS regulation.

We do not know how a decreased NADH/NAD+ ratio induces apoptosis in the cells with high LIP expression. NAD+ is an essential co-factor for metabolic reactions and a substrate for reactions in cell signalling pathways, including sirtuins and poly-adenosine ribose-polymerase (PARP) (Canto et al., 2015; Canto et al., 2013). Increased NAD+ levels rather protect from apoptosis through sirtuin mediated mechanisms (Verdin et al., 2010). Recently, it was shown that NADH is able to bind to the apoptosis inducing factor (AIF), inducing its dimerization and thereby maintaining the AIF-dimers in the mitochondria preventing apoptosis (Brosey et al., 2016). Low levels of NADH results in AIF monomerization and the AIF monomers can leave the mitochondria and translocate into the nucleus to induce apoptosis (Sevrioukova, 2011). However, the induced MAS activity would remove NADH from the cytosol but not from the mitochondria suggesting a different, yet to be identified mechanism. Although we could not clarify the mechanism of apoptosis induction by 2-DG treatment in cells with high LIP expression, we demonstrate in this manuscript that the activation of the MAS may become an “Achilles heel” for cancer cells upon glucose restriction. We show that upon glucose starvation or inhibition of glycolysis a simultaneously active MAS creates low NADH/NAD+ ratios that are toxic to the cells.

New attempts in cancer therapy try to exploit cancer cell specific metabolic dependencies to specifically kill cancer cells. So far, cancer therapy with 2-DG is only tested for a few specific cancer types or in combination with other chemotherapeutic drugs (Zhang et al., 2014). Our study suggests that high LIP expression may be used as a biomarker for effectiveness of 2-DG or other glycolysis inhibiting drugs in cancer treatment. Further work is needed to evaluate the predictive power of LIP expression for 2-DG treatment success. Furthermore, we identified the NADH/NAD+ ratio as an important mediator of 2-DG toxicity *in vitro*. More experiments are required to evaluate whether 2-DG lowers the NADH/NAD+ ratio in tumours or whether artificial oxidation of NADH with small compounds (e.g. duroquinone) will make cancer cells sensitive to 2-DG treatment duroquinone (Gui et al., 2016).

Taken together, we found that LIP renders cells sensitive to 2-DG treatment by stimulating the MAS and its use of cytoplasmic NADH. We describe that the consequently low NADH/NAD+ ratio is an important mediator of 2-DG induced cell death in triple negative breast (TNBC) cancer cells. Furthermore, we suggest a new model for 2-DG sensitivity in which a low NADH/NAD+ ratio mediated by high LIP/high MAS or other hypothetical mechanisms drive cells into apoptosis.

## Acknowledgements

At the UMCG we thank Stefan Juranek and Floris Foijer for supervising the CRISPR/CAS9 facility where BT20 *Cebpb* knockout cells were generated and Hilde Jalving for providing the BT549, MDA-MB-231 and ZR-75-1 breast cancer cell lines. We like to thank Christine Müller for advice on writing the manuscript. T.A. and G.K. were supported by the Dutch Cancer Society (KWF #10080) and C.M. and G.H. were supported by the Deutsche Krebshilfe through grant (DKH #612100) through grants to C.FC.

## Authorship contributions

T.A. designed and performed the research, and collected and analysed the data; H.R.Z., G.K, M.A.Z. and M.R. performed research and collected data; G.H. and C.F.C. designed research and supervised the project; T.A. and C.F.C. wrote the manuscript.

## Methods

### Cell culture

BT20 cells, MDA-MB-231 and all immortalized MEF cell lines were culture in high glucose DMEM supplemented with 10% FBS, 10mM HEPES, 1mM Sodium Pyruvate and 100U/ml Penicillin Streptomycin. BT549, ZR-75-1, T47D and MCF7 breast cancer cells were maintained in RPMI1640 medium supplemented with 10% FBS, 25mM HEPES, 1mM Sodium Pyruvate and 100U/ml Penicillin/Streptomycin. C/EBP□ ko MEF were described before (Zidek et al., 2015).

### DNA constructs

Plasmids containing rat C/EBPβ-LAP, rat C/EBPβ-LIP and human C/EBPβ-LIP and were described before (Zidek et al., 2015). For CRISPR/Cas9 mediated knock out of both C/EBPβ isoforms, guide RNA squences (LAP: 5’-GAGTGGCCAACTTCTACTACG-3’, LIP: 5’-GCGCTTACCTCGGCTACCAGG-3’) were cloned into pSpCas9(BB)-2A-puro (PX459) v2.0 (http://www.addgene.org/62988/).

### Transfection

Immortalized MEFs were transfected with an empty, rat C/EBP□-LIP or -LAP containing pcDNA3 or pSV2Zeo vector by using FugeneHD (Promega) according to the manufactures protocol. For stable overexpression, C/EBP□□ko MEFs were treated with 0.2 mg/ml Zeocin (Invitrogen). To maintain the expression cells were culture with 0.1 mg/ml Zeocin in the medium. T47D cells and MCF7 cells were transfected with empty or human LIP-containing pcDNA3.1 via Fugene HD (Promega) using the manufactures protocol. For stable expression, MCF7 cells were selected with 0.8 mg/ml, T47D with 0.4mg/ml G418. For CRISPR/Cas9 mediated knock out of both C/EBPβ isoforms, BT20 cells cells were transfected with Fugene HD (Promega) according to the manufactures protocol and selected with puromycine (1 µg/ml). After the selection, clones were grown out and C/EBPβ level were analysed by western blot.

### Proliferation assays

To determine the proliferation and survival of cancer cells, relative cell numbers were measured using the CellTiter-Fluor™ Cell Viability Assay (Promega) after 3 days of treatment. Measurements were performed according to the suppliers manual.

### Metabolic flux analysis

Metabolic flux analysis was performed using a Seahorse XF96 Extracellular Flux analyser (Agilent Bioscience). 1.5×10^4^ EV, LAP or LIP overexpressing C/EBPβ ko MEFs were seeded 4h before the assay. Assays were performed according to the manufactures protocol. Injected drugs were oligomycin (2,5 µM) for blockage of ATP related respiration, dinitrophenol (50 µM) to uncouple the mitochondrial membrane (maximum respiration) and 2-DG (100 mM) for inhibition of glycolysis. 3×10^4^ EV and LIP overexpressing T47D cells were seeded 16 hours before the experiments. Assays were performed according to the manufactures protocol. Injected drugs were UK5099 (5 µM) for blockage of mitochondrial pyruvate transporter, BPTES (3 µM) for blockage of glutaminase, etomoxir (40 µM) for blockage of fatty acid transport into the mitochondrion, 2-DG (100 mM) for inhibition of glycolysis, Rotenone (4 µM) blockage of complex 1 and oligomycin (2,5 µM) for blockage of ATP related respiration. To measure the activity of the MAS, cells were permeabilized by injection of saponin (25 µg/ml) and the substrates of the MAS (Glutamate (1 mM) and Malate (2 mM)) were added separately by single injections.

### Luciferase based assays

NADH, NAD+, ATP and ADP level were distinguished using luciferase based assays. 24 hours before the assay, 7500 cells per well were seeded in a 96-well plate. Experiments were performed according to manufactures protocols (NADH/NAD+: Promega, G9071; ATP/ADP: Biovision, K255-200). For detection, a GloMax-Multi Detection System (Promega) was used. Caspase3/7 activity was measured 3 days after the treatment with a commercial available kit (Caspase-Glo 3/7 Assay, Promega).

### Immunoblot analysis

Cells and tissues were lysed using RIPA buffer. Equal amounts of protein were separated via SDS-PAGE and transferred to a PVDF membrane using Trans-Blot Turbo System (Bio-rad). The following antibodies were used for detection: C/EBPβ (E299) from Abcam, □-Tubulin (GT114) from GeneTex and β-actin (clone C4) (#691001) from MP Biomedicals. For detection, HRP-conjugated secondary antibodies (Amersham Life Technologies) were used. The signals were visualized by chemiluminescence (ECL, Amersham Life Technologies) using ImageQuant LAS 4000 mini imaging machine (GE Healthcare Bioscience AB) and the supplied software was used for the quantification of the bands.

### Statistical analysis

IC50 and r2 values were calculated using a non/linear fitting algorithm (log[inhibitor] vs response - variable slope (four parameters)) in Graphpad Prism.

---

**Supplementary Figure 1.**
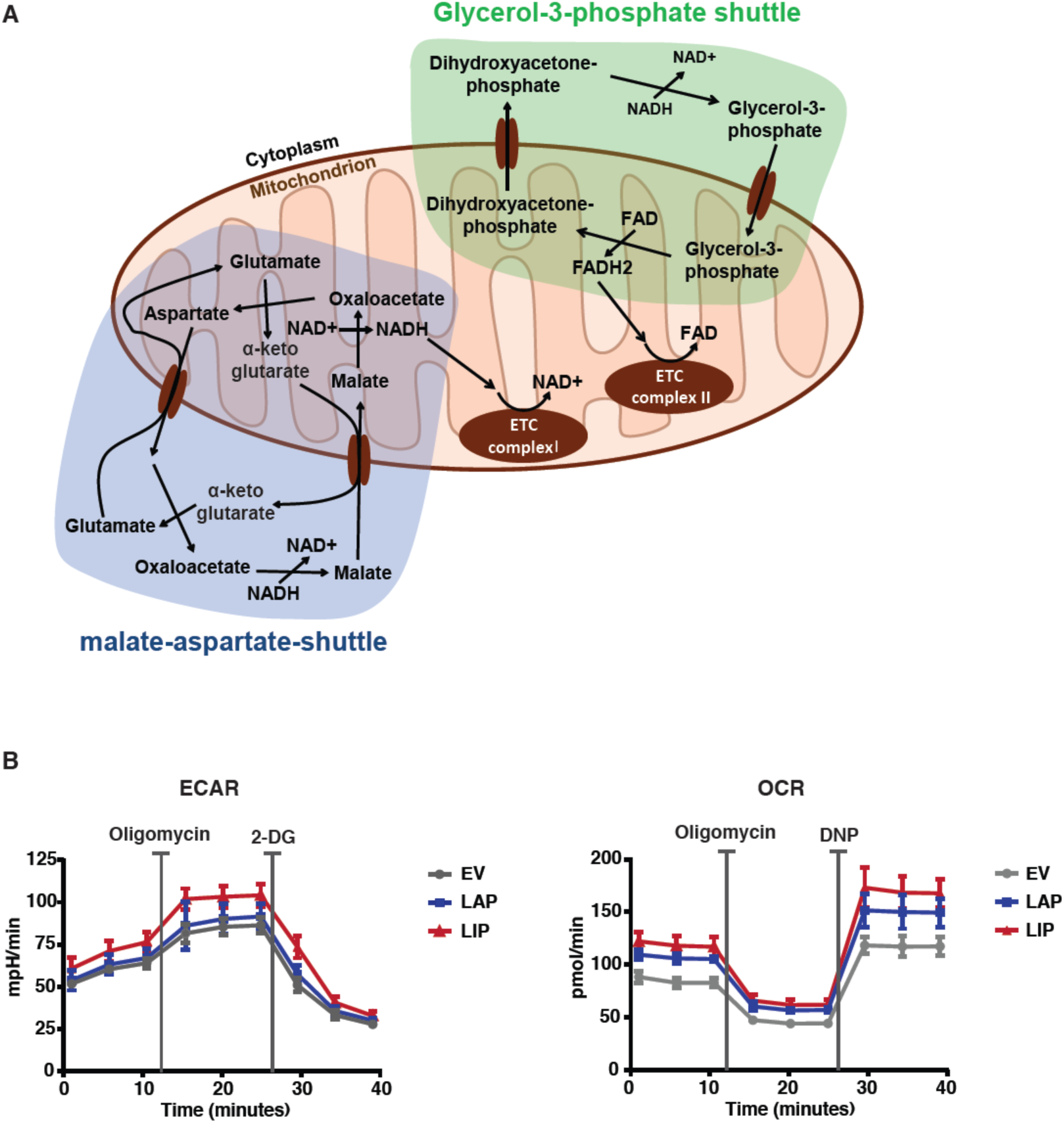
(**A**) Schematic representation of malate-aspartate-shuttle (MAS) and glycerol-3-phosphate-shuttle (GPS). (**B**) ECAR OCR of *Cebpb*-ko fibroblasts transfected with expression vectors for LAP, LIP or empty vector (EV) control (n=6).

**Supplementary Figure 2.**
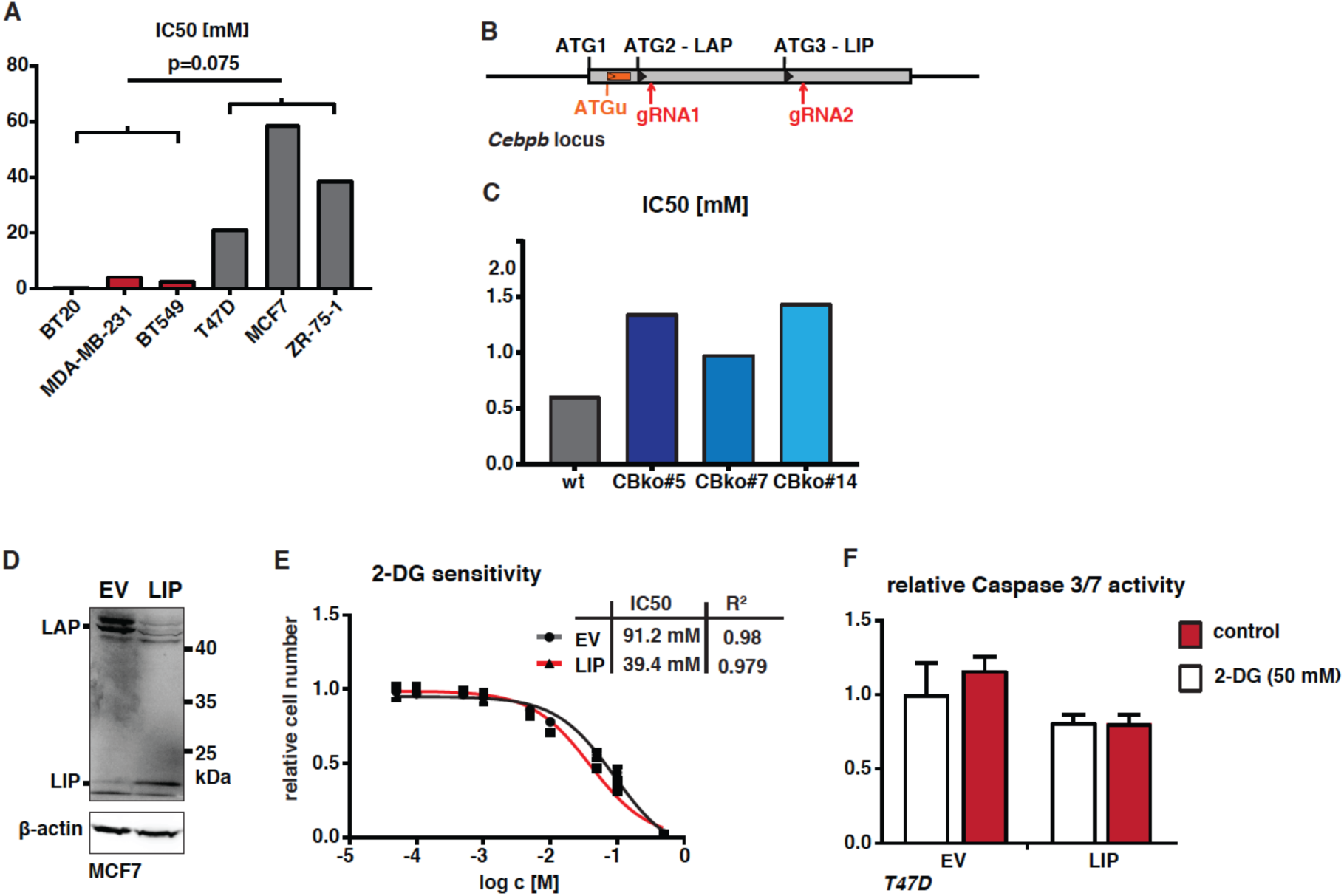
(**A**) Bar graph representation of IC50 values of 2-DG treatment of the TNBC BT20, MDA-MB-231, BT549, and Luminal A MCF7, T47D, ZR-75-1 breast cancer cell lines (related to Figure 2B). (**B**) Schematic representation of *Cebpb* locus and the CRISPR/Cas9 based knockout strategy. The arrows mark the guide-RNAs used. (**C**) Bar graph representation of IC50 values of 2-DG treatment of wt BT20 cells and three clones of CRIPR/Cas9 derived *Cebpb*-ko BT20 cells (related to Figure 2E). (**D**) Immunoblot showing expression of LAP, LIP and β-actin as loading control in MCF7 cells transfected with expression vectors for LIP or empty vector (EV) control. (**E**) Dose-response-curve MCF7 cells expressing LIP or empty vector (EV) control after 3 days of 2-DG treatment (n=5). (**F**) Relative Caspase 3/7 activity of EV control and LIP overexpressing T47D cells after 3 days of 2-DG treatment (n=5). IC50 values of dose-response are shown with statistical analysis in R^2^.

**Supplementary Figure 3.**
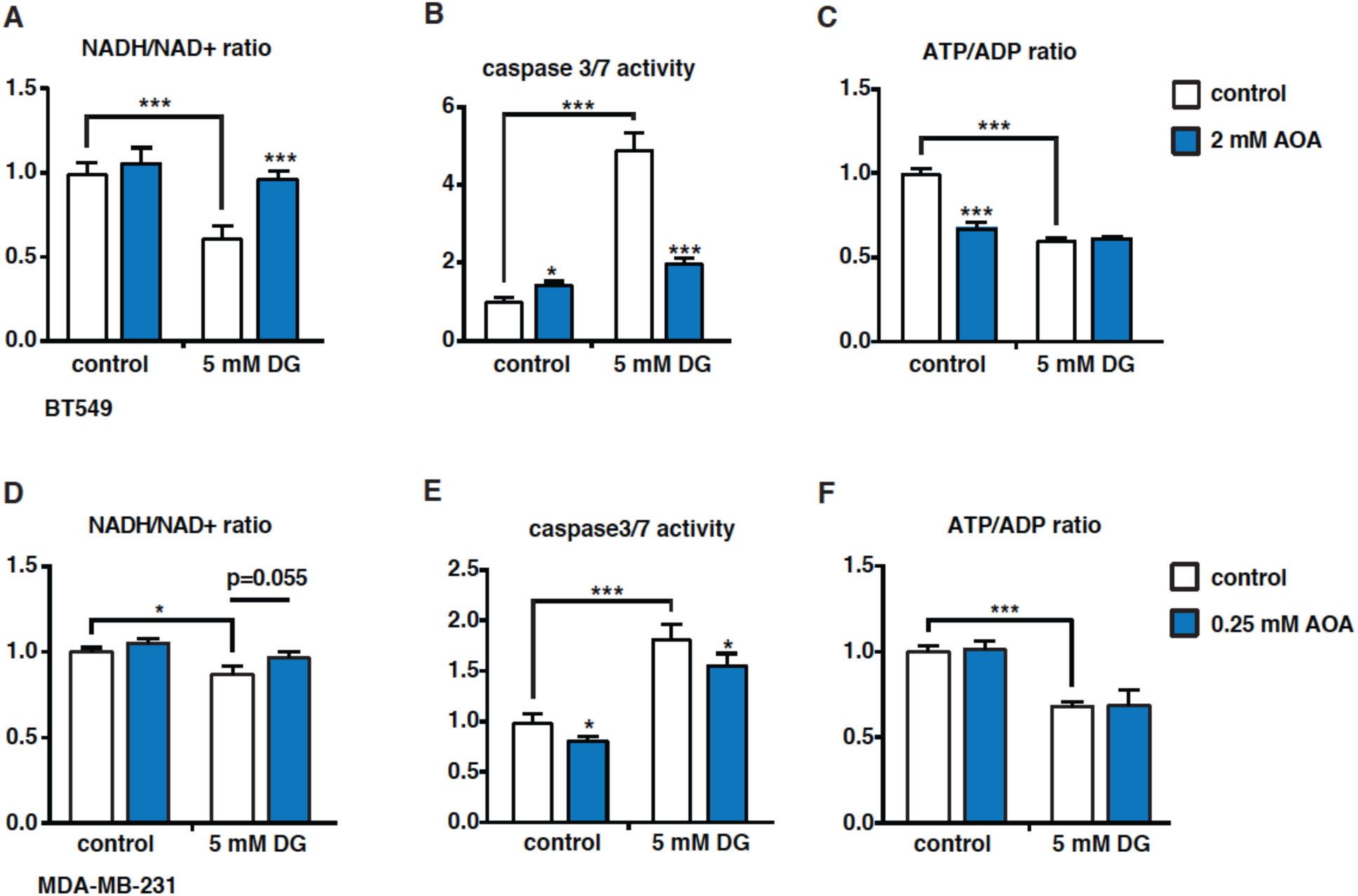
**(A**) Relative NADH/NAD+ ratios in BT549 cells after 1 day of treatment with solvent (control), 2-DG, AOA, or 2-DG and AOA (n=4). (**B**) Relative caspase3/7 activity of BT549 cells after 3 days of treatment with solvent, 2-DG, AOA, or 2-DG and AOA (n=4). (**C**) Relative ATP/ADP ratios in BT549 cells after 1 day of treatment with solvent, 2-DG, AOA, or 2-DG and AOA (n=4). (**D**) Relative NADH/NAD+ ratios in MDA-MB-231 cells after 1 day of treatment with solvent, 2-DG, AOA, or 2-DG and AOA (n=3). (**E**) Relative ATP/ADP ratios in MDA-MB-231 cells after 1 day of treatment with solvent, 2-DG, AOA, or 2-DG and AOA (n=3). (**F**) Relative caspase3/7 activity in MDA-MB-231 cells after 3 days of treatment with solvent, 2-DG, AOA, or 2-DG and AOA (n=4). (**G**) Relative NADH/NAD+ ratios T47D cells expressing LIP or empty vector (EV) control after 1 day of treatment with solvent, 2-DG, AOA or 2-DG and AOA (n=4). Statistical differences were analysed by Student’s t-tests. Error bars represent SD, * p<0.05, *** p<0. 001.

## References

Ackermann, T., Hartleben, G., Muller, C., Mastrobuoni, G., Groth, M., Sterken, B.A., Zaini, M.A., Youssef, S.A., Zuidhof, H.R., Krauss, S.R., et al. (2019). C/EBPbeta-LIP induces cancer-type metabolic reprogramming by regulating the let-7/LIN28B circuit in mice. Commun Biol 2, 208.

Begay, V., Smink, J.J., Loddenkemper, C., Zimmermann, K., Rudolph, C., Scheller, M., Steinemann, D., Leser, U., Schlegelberger, B., Stein, H., et al. (2015). Deregulation of the endogenous C/EBPbeta LIP isoform predisposes to tumorigenesis. J Mol Med (Berl) 93, 39–49.

Birsoy, K., Wang, T., Chen, W.W., Freinkman, E., Abu-Remaileh, M., and Sabatini, D.M. (2015). An Essential Role of the Mitochondrial Electron Transport Chain in Cell Proliferation Is to Enable Aspartate Synthesis. Cell 162, 540–551.

Borst, p. (2020). The malate-aspartate shuttle (Borst cycle): How it started and developed into a major metabolic pathway. IUBMB Life.

Brosey, C.A., Ho, C., Long, W.Z., Singh, S., Burnett, K., Hura, G.L., Nix, J.C., Bowman, G.R., Ellenberger, T., and Tainer, J.A. (2016). Defining NADH-Driven Allostery Regulating Apoptosis-Inducing Factor. Structure 24, 2067–2079.

Calkhoven, C.F., Bouwman, P.R., Snippe, L., and Ab, G. (1994). Translation start site multiplicity of the CCAAT/enhancer binding protein alpha mRNA is dictated by a small 5’ open reading frame. Nucleic Acids Res 22, 5540–5547.

Calkhoven, C.F., Muller, C., and Leutz, A. (2000). Translational control of C/EBPalpha and C/EBPbeta isoform expression. Genes Dev 14, 1920–1932.

Canto, C., Menzies, K.J., and Auwerx, J. (2015). NAD(+) Metabolism and the Control of Energy Homeostasis: A Balancing Act between Mitochondria and the Nucleus. Cell Metab 22, 31–53.

Canto, C., Sauve, A.A., and Bai, p. (2013). Crosstalk between poly(ADP-ribose) polymerase and sirtuin enzymes. Mol Aspects Med 34, 1168–1201.

DeBerardinis, R.J., and Chandel, N.S. (2016). Fundamentals of cancer metabolism. Sci Adv 2, e1600200.

DeBerardinis, R.J., Lum, J.J., Hatzivassiliou, G., and Thompson, C.B. (2008). The biology of cancer: metabolic reprogramming fuels cell growth and proliferation. Cell Metab 7, 11–20.

Descombes, P., and Schibler, U. (1991). A liver-enriched transcriptional activator protein, LAP, and a transcriptional inhibitory protein, LIP, are translated from the same mRNA. Cell 67, 569–579.

Greenhouse, W.V., and Lehninger, A.L. (1976). Occurrence of the malate-aspartate shuttle in various tumor types. Cancer Res 36, 1392–1396.

Gui, D.Y., Sullivan, L.B., Luengo, A., Hosios, A.M., Bush, L.N., Gitego, N., Davidson, S.M., Freinkman, E., Thomas, C.J., and Vander Heiden, M.G. (2016). Environment Dictates Dependence on Mitochondrial Complex I for NAD+ and Aspartate Production and Determines Cancer Cell Sensitivity to Metformin. Cell Metab 24, 716–727.

Hanahan, D., and Weinberg, R.A. (2011). Hallmarks of cancer: the next generation. Cell 144, 646–674.

Hanse, E.A., Ruan, C., Kachman, M., Wang, D., Lowman, X.H., and Kelekar, A. (2017). Cytosolic malate dehydrogenase activity helps support glycolysis in actively proliferating cells and cancer. Oncogene 36, 3915–3924.

Hung, Y.P., Albeck, J.G., Tantama, M., and Yellen, G. (2011). Imaging cytosolic NADH-NAD(+) redox state with a genetically encoded fluorescent biosensor. Cell Metab 14, 545–554.

Jundt, F., Raetzel, N., Müller, C., Calkhoven, C.F., Kley, K., Mathas, S., Lietz, A., Leutz, A., and Dorken, B. (2005). A rapamycin derivative (everolimus) controls proliferation through down-regulation of truncated CCAAT enhancer binding protein {beta} and NF-{kappa}B activity in Hodgkin and anaplastic large cell lymphomas. Blood 106, 1801–1807.

Koppenol, W.H., Bounds, P.L., and Dang, C.V. (2011). Otto Warburg’s contributions to current concepts of cancer metabolism. Nat Rev Cancer 11, 325–337.

Millward, C.A., Heaney, J.D., Sinasac, D.S., Chu, E.C., Bederman, I.R., Gilge, D.A., Previs, S.F., and Croniger, C.M. (2007). Mice with a deletion in the gene for CCAAT/enhancer-binding protein beta are protected against diet-induced obesity. Diabetes 56, 161–167.

Müller, M., Zidek, L.M., Ackermann, T., de Jong, T., Liu, P., Kliche, V., Zaini, M.A., Kortman, G., Harkema, L., Verbeek, D.S., et al. (2018). Reduced expression of C/EBPβ-LIP extends health-and lifespan in mice. bioRxiv.

Pavlova, N.N., and Thompson, C.B. (2016). The Emerging Hallmarks of Cancer Metabolism. Cell Metab 23, 27–47.

Quintanilla-Martinez, L., Pittaluga, S., Miething, C., Klier, M., Rudelius, M., Davies-Hill, T., Anastasov, N., Martinez, A., Vivero, A., Duyster, J., et al. (2006). NPM-ALK-dependent expression of the transcription factor CCAAT/enhancer binding protein beta in ALK-positive anaplastic large cell lymphoma. Blood 108, 2029–2036.

Rask, K., Thorn, M., Ponten, F., Kraaz, W., Sundfeldt, K., Hedin, L., and Enerback, S. (2000). Increased expression of the transcription factors CCAAT-enhancer binding protein-beta (C/EBBeta) and C/EBzeta (CHOP) correlate with invasiveness of human colorectal cancer. Int J Cancer 86, 337–343.

Rej, R. (1979). Measurement of aspartate aminotransferase activity: effects of oxamate. Clin Chem 25, 555–559.

Sevrioukova, I.F. (2011). Apoptosis-inducing factor: structure, function, and redox regulation. Antioxid Redox Signal 14, 2545–2579.

Sundfeldt, K., Ivarsson, K., Carlsson, M., Enerback, S., Janson, P.O., Brannstrom, M., and Hedin, L. (1999). The expression of CCAAT/enhancer binding protein (C/EBP) in the human ovary in vivo: specific increase in C/EBPbeta during epithelial tumour progression. Br J Cancer 79, 1240–1248.

Tajan, M., Hock, A.K., Blagih, J., Robertson, N.A., Labuschagne, C.F., Kruiswijk, F., Humpton, T.J., Adams, P.D., and Vousden, K.H. (2018). A Role for p53 in the Adaptation to Glutamine Starvation through the Expression of SLC1A3. Cell Metab 28, 721–736 e726.

Thornburg, J.M., Nelson, K.K., Clem, B.F., Lane, A.N., Arumugam, S., Simmons, A., Eaton, J.W., Telang, S., and Chesney, J. (2008). Targeting aspartate aminotransferase in breast cancer. Breast Cancer Res 10, R84.

Vander Heiden, M.G., Cantley, L.C., and Thompson, C.B. (2009). Understanding the Warburg effect: the metabolic requirements of cell proliferation. Science 324, 1029–1033.

Vander Heiden, M.G., and DeBerardinis, R.J. (2017). Understanding the Intersections between Metabolism and Cancer Biology. Cell 168, 657–669.

Verdin, E., Hirschey, M.D., Finley, L.W., and Haigis, M.C. (2010). Sirtuin regulation of mitochondria: energy production, apoptosis, and signaling. Trends Biochem Sci 35, 669–675.

Wang, Y.P., Zhou, W., Wang, J., Huang, X., Zuo, Y., Wang, T.S., Gao, X., Xu, Y.Y., Zou, S.W., Liu, Y.B., et al. (2016). Arginine Methylation of MDH1 by CARM1 Inhibits Glutamine Metabolism and Suppresses Pancreatic Cancer. Mol Cell 64, 673–687.

Zahnow, C.A., Cardiff, R.D., Laucirica, R., Medina, D., and Rosen, J.M. (2001). A role for CCAAT/enhancer binding protein beta-liver-enriched inhibitory protein in mammary epithelial cell proliferation. Cancer Res 61, 261–269.

Zahnow, C.A., Younes, P., Laucirica, R., and Rosen, J.M. (1997). Overexpression of C/EBPbeta-LIP, a naturally occurring, dominant-negative transcription factor, in human breast cancer. J Natl Cancer Inst 89, 1887–1891.

Zhang, D., Li, J., Wang, F., Hu, J., Wang, S., and Sun, Y. (2014). 2-Deoxy-D-glucose targeting of glucose metabolism in cancer cells as a potential therapy. Cancer Lett 355, 176–183.

Zidek, L.M., Ackermann, T., Hartleben, G., Eichwald, S., Kortman, G., Kiehntopf, M., Leutz, A., Sonenberg, N., Wang, Z.Q., von Maltzahn, J., et al. (2015). Deficiency in mTORC1-controlled C/EBPbeta-mRNA translation improves metabolic health in mice. EMBO Rep 16, 1022–1036.

